# Phytochemical characterization and antimicrobial properties of *Azadirachtin indica* against selected bacterial human pathogens

**DOI:** 10.1101/2024.10.29.620911

**Authors:** Idorenyin Idorenyin Eka, Idongesit Udo Wilson, Aniefon Alphonsus Ibuot

## Abstract

The study was conducted to assess the phytochemical and antibacterial potentials of *Azadirechtin indica* against *Salmonella typhi, Staphylococcus aureus* and *Pseudomonas aeruginosa* using microbiological techniques. Ethanol and water were used to extract the active components from the *Azadirechtin indica* leaves. *Salmonella typhi* showed range of inhibition zone from 11mm to 16mm at 50% to 90% concentrations of the plant ethanolic extract. *Staphylococcus aureus* had a range of inhibition zone from 10mm to 13mm at 50% to 90 % concentrations of the plant ethanolic extract and *Pseudomonas aeruginosa* had a range of inhibition zone from 10mm to 12mm at 50% to 90% concentrations of the plant ethanolic extract. Ethanol extract showed greater antibacterial activity against the test organisms when compared with the aqueous extract. Preliminary screening of phytochemicals in the leaf extract of *Azadirechtin indica* revealed the presence of alkaloids (7.376mg/100g), saponins (12.041mg/100g), flavonoids (2.641mg/100g), tannins (3.431mg/100g) and cyannogenic glycosides (0.952 mg/100g). This result provides scientific support for *Azadirechtin indica* to be further investigated and used as drug.

## 1.0 Introduction

All through human history Medicinal plants have been used in the treatments of diseases and ailments all through human history. Medicinal plants are plant that contents several chemical compounds which are reported to play significant role in human biological function and also prevent microorganisms and insects from causing harm. Medicinal plants have been reported to comprise a natural reservoir for medicines in the world [1]. They are central in traditional medicine and are the core therapeutic.

Several medicinal plants are made of different chemical compounds such as flavonoids, steriods, alkaloids, saponins, anthraquinones, glycosides and tannins [2]. About 12,000 out of the chemical compounds identified in medicinal plants have been discovered, a fraction which is considered to be very insignificant, less than the 10% of the total chemical compounds contain in medicinal plants. Several plants have been discovered to contain several metabolites which have antimicrobial properties. Although most of these medicinal plants are currently being used especially in developing nations in the treatment of diseases, a lot of the medicinal plants are yet to be identified and the full health benefits of these plants are yet to be unraveled [2].

Before modern medicine was introduced, people relied on medicinal plants (roots, leaves, bark) for their health wellbeing. The challenge was the dosage of administration and preparations which may have been dangerous to human health. The medicinal plants have substantial active ingredients which could cure or give relief to health issues. Several of these active ingredients are useful drugs and some have been modified to form synthetic drugs [1].

*Azadirachta indica* is abundantly present in the tropical countries of the world and is well known for its insecticidal and various types of antimicrobial properties. Almost every part of this plant tree has been reported to possess a several pharmacological properties. The medicinal and insecticidal properties of different parts of *Azadirachta indica* tree have been well documented. The leaves have antimicrobial properties and are traditionally being used as curative agent against certain fungal and bacterial diseases [3]. This work is therefore aimed at evaluating the antimicrobial and phytochemical properties of *Azadirachtin indica*

## 2.0 Methods

### 2.1. Sample collection and preparation

Fresh leaves of *Azadirachtin indica* plant were collected from the botanical garden. The plant sample was stored in a sterile polythene bag and taken to the microbiology laboratory in Akwa Ibom State polytechnic, for analysis. The leaves were washed thoroughly with distilled water and kept to dry. They were sliced into smaller sizes and air-dried at room temperature to remove the moisture content.

The dried sample were crushed into powdery form using a sterile blender, a fine powder was obtained and stored in an air-tight container.

### 2.2 Extraction of ethanolic and aqueous extract

To prepare ethanolic and aqueous extract of the plant sample, 100 g of the grounded leaves were weighed into a flask containing 200 ml of 75 % ethanol and soaked instantly at an interval of 20 minutes for three days for proper extraction. And was filtered using Whatman filter paper, the filtrate was transferred into a beaker and placed in a water bath regulated at temperature of 50 ^0^C for dryness through evaporation. The same procedure was carried out for aqueous extract; it was soaked for 24 hours before filtering with Whatman filter paper. The dried extract (paste) was carefully transferred into a sample bottle sealed immediately and stored in the refrigerator at 4 °C for further analysis of antimicrobial activity.

### 2.3 Morphological and Biochemical identification of bacterial isolates

*Salmonella typhi, Staphylococcus aereus* and *Pseudomonas aeruginosa* were cultured in the laboratory. Adequate inspections were carried out for all the isolates, the cultural characteristics were noted. The biochemical characterization as described by Cheesbrough [4] were carried out which included; catalase, coagulase, oxidase, indole, citrate, urease, motility, spores, methyl red-voges proskaur test and carbohydrate fermentation.

### 2.4 Preparation of Antibacterial Sensitivity disc and inoculation of test bacteria

The residue obtained from the evaporated extract was dissolved with ethanol for ethanolic extract and distilled water for aqueous extract respectively and was subjected to different concentrations ranging from 10 mg/ml, 7.5 mg/ml and 5 mg/ml and Perforated Whatman paper No 1 of about 6 mm in diameter sterilized in hot air oven and allowed to cooling, was soaked in different concentrations of the extract, removed and was allowed for few minutes to dry. According to NCCLS, [6] molten agar was poured into sterile petri dishes and allowed to solidify. A sterile wire loop was used to pick a loopful of the test isolates (*Salmonella typhi* and *Staphylococcus aureus*) inoculated differently and uniformly on the labeled solidified agar plates. Each of the prepared discs was placed equidistant in all the plates based on the concentrations. The plates were wrapped with aluminum foil and invertedly incubated at 37^0^C for 18-24 hours for detection of clear zones of inhibition. The inhibitory zones were interpreted as sensitive, intermediary or resistant, using the interpretation chart of zone size of Kirby-Bauer sensitivity test method described by Prescott et al., [5]. Interpretations of the result was done using inhibiting zone sizes, 18 mm and above were considered sensitive, 13-17 mm intermediate and 13 mm below as resistant according to NCCLS [6].

### 2.5 Phytochemical Screening of the plant sample

The powdered form of the leave sample was then subjected to phytochemical analysis to determine the bioactive components present in the leave sample. The bioactive component includes Alkaloids, flavonoids, saponins, Tanins and Cyanogenic glycosides was carried out using the method of Edeoga *et al* [7] and Oluduro [8]

## 3.0 Results

### 3.1. Morphological and biochemical characteristics of bacterial isolates

The phenotypic and biochemical characteristics of bacterial isolates were carried out for identification as shown in tables 1 and 2.

**Table 1:** Biochemical characteristics of bacterial isolates.

| Isolates | Gram stain | Spore | Catalase | Coagulase | Motility | Citrate | Urease | Methyl Red | V.P | Indole | Oxidase | Glucose | Lactose | Maltose | Sucrose |
| --- | --- | --- | --- | --- | --- | --- | --- | --- | --- | --- | --- | --- | --- | --- | --- |
| 1 | + | - | + | + | - | + | + | + | + | - | - | AG | AG | AG | AG |
| 2 | - | - | + | - | + | - | - | + | - | - | - | AG | AO | AG | AO |
| 3 | - | - | + | - | + | + | - | - | - | - | + | AO | AO | AG | AO |

+ positive
-negative
A- Acid production
G- Gas formation
O- No reaction

**Table 2:** Phenotypic characteristics of bacterial isolates.

| Isolates | Cultural and morphological characteristics | Cell shape | Probable Bacteria |
| --- | --- | --- | --- |
| 1 | Circular, pink, raised, entire, opaque | Cocci in cluster | *Staphylococcus aureus* |
| 2 | Irregular, dark, umbonate, undulate, opaque | Rods | *Salmonella typhi* |
| 3 | Irregular, light green, flat, dentate, translucent | Rods in short chains | *Pseudomonas aeruginosa* |

### 3.2. Antibiogram of Bacterial Isolates on different concentrations of plant extract

Antibacterial effect of *Azadirachtin indica* leaves extract on the four tests isolates was carried out using agar disc diffusion method (Table 3).

**Table 3:** Antibiogram of test bacterial isolates to plant extracts.

| Isolates | Ethanolic extract |  |  | Aqueous extract |  |  |
| --- | --- | --- | --- | --- | --- | --- |
|  | 90% | 70% | 50% | 90% | 70% | 50% |
| *Staphylococcus aureus* | 13mm | 11mm | 10 mm | - | 7mm | 8mm |
| *Salmonella typhi* | 11mm | 12mm | 16mm | - | - | 9mm |
| *Pseudomonas aeruginosa* | - | 10mm | 12mm | - | - | - |

### 3.3. Phytochemical analysis of *Azadirachtin indica*

The phytochemical screening of *Azadirachtin indica* leaves showed bioactive components which include; alkaloids, flavonoids, saponins, tannins and cyanogenic glycosides (Table 4).

**Table 4:** Phytochemical screening of *Azadirachtin indica* leaves.

| Parameters | Concentration { Mg/100g } |
| --- | --- |
| Alkaloids | 7.376 ± 0.411 |
| Saponins | 12.041 ± 0.212 |
| Flavonoids | 2.601 ± 0.109 |
| Tannins | 3.431 ± 0.198 |
| Cyanogenic glycosides | 0.952 ± 0.065 |

## 4.0 Discussions

Infectious diseases are a major cause of mortality and morbidity worldwide, currently the ongoing battle against bacteria and their evolving resistance is of serious concern. Signaling the right time to discover new plant –based antimicrobial agent without the tendency to be resisted by pathogens [9].Several investigations conducted in many countries, confirms the presence of active compounds in medicinal plants [10,11]. Vijayan *et al*., [12] reports that more than 80% of the world depends on herbal medicine. Plant parts have been reported to serve as sources of new medicinal drugs [13]. Currently, there has been a significant switch towards herbal medicines due to resistance of pathogens to antibiotics and irreversible reactions caused by of modern drugs. The plant resources and herbal reserves tend to be exhausted due to urbanization and exploitation of plants reservoir [3]. In the current times of drug discovery, several plant products are assessed on the basis of their antimicrobial significance and traditional uses.

In many areas of antibacterial discovery, *A. indica* has been reported to be effective against several human pathogens. Overall, a large number of studies have been published in the last decade on this topic, especially as they relate to the ever-growing number of antibiotic-resistant organisms. Hence, several bacterial species are found on the long list of antibiotic resistant threats which is a great concern, including *S. aureus* and *P. aeruginosa* [14]. Several researchers have tested the antibacterial effect of *A. indica* against these species. Garg *et al*. [15] reported that methanol and chloroform *A. indica* extracts exhibited more antimicrobial effect against *S. aurues* and *P. aeruginosa* better than other plant extracts and even better than several different antibiotics. More specifically, it has also been demonstrated that the MIC of limonoid compounds isolated from *A. indica* seeds had between 32 μg/ml and 128 μg/ml concentration against *P. aeruginosa* and some opportunistic skin pathogens, *Staphylococcus epidermidis* [16]. Several studies have showed that an aqueous *A. indica* leaf extract was used to make alginate fibers for wound dressings [17], a nanofibrous mat embedded with *A. indica* leaf extract [18], and a topical gel containing a methanolic *A. indica* extract all had inhibitory effect on *S. aureus* growth [19]. Killi *et al*., [20] reported that polyesteramide synthesized from *A. indica* oil to produce a nanofibrous mat, the incorporation of *A. indica* into this wound treatment method resulted in increased tissue regeneration in rats, as compared to the control commercial cream [20].These reports further confirms the antimicrobial strength of *A. indica* against *S. aureus, Salmonella typhi* and *Pseudomanas aeruginosa* as reported in this study.

## 5.0 Conclusion

This study reveals that the ethanolic extracts of Azadirechtin indica possess and exhibit antibacterial effect against some pathogenic bacteria. However, further investigation of the plant parts is needed for more scientific confirmation and subsequent use of the plant as drug for treatment of diseases caused by the test pathogens

## Acknowledgements

The article was sponsored by TETFund research grant, Nigeria

## Disclosure of conflict of interest

The authors declare no conflict of interest

## Notes

### Competing Interest Statement

The authors have declared no competing interest.

